# Lysophosphatidic acid controls autotaxin binding to β1-integrin : impact on disease progression in murine collagen-induced arthritis

**DOI:** 10.1101/2025.04.04.646839

**Authors:** Mathias Emery, Cinthia-Nayeli Cuero, Lamia Bouazza, Candice Internicola, François Duboeuf, Patricia Cerecero Aguirre, Andrew A. McCarthy, Irma Machuca-Gayet, Raphaël Leblanc, Olivier Peyruchaud

**Affiliations:** European Molecular Biology Laboratory, EMBL Grenoble, 71 avenue des Martyrs, CS 9018, 38042 Grenoble, France; Universite Claude Bernard Lyon 1, INSERM, UMR 1033, LYOS, Lyon, France; Universidad Autonoma del Estado de México, Toluca, México; Universite Claude Bernard Lyon 1, INSERM, CNRS, UMR 1033, LYOS, Lyon, France; Centre de Cancérologie de Marseille, Marseille, France

## Abstract

Autotaxin (ATX) is a lysophospholipase D (lysoPLD) serving as both a lysophosphatidic acid (LPA)-producing enzyme and a LPA docking molecule. ATX binds to the cell surface via interaction with adhesive molecules, including β1-integrin, potentially facilitating LPA access to its specific G protein-coupled receptors. However, the precise protein-protein interaction sequences and their biological implications remain unknown. Here, we identify the interaction domains between ATX and β1-integrin and generate specific blocking antibodies allowing to demonstrate that ATX-β1 binding domain involves a cryptic epitope unmasked by LPA docking. In addition, whereas anti-ATX antibodies do not inhibit the lysoPLD activity, immunological neutralization of the ATX-β1 integrin binding site reduces arthritis development in a collagen-induced arthritis model. These findings offer novel insights into the molecular mechanisms governing ATX functions, which, in addition to its enzymatic activity, requires cell surface binding. These findings suggest that ATX binding domains could be targeted for novel therapeutic approaches.

## INTRODUCTION

Autotaxin (ATX, ENPP2) is a secreted member of the ectonucleotide pyrophosphatase/phosphodiesterase (ENPP) family, characterized with a unique lysophospholipase D (LysoPLD) activity^1^. This enzymatic activity enables the hydrolysis of lysophosphatidylcholine (LPC) and other extracellular lysophospholipids to produce the bioactive lipid lysophosphatidic acid (LPA), which subsequently activates a set of six G protein-coupled receptors (LPAR) ^2^. Both ATX and LPA are physiological components found of biological liquids, including saliva, blood, urine, seminal fluid, and cerebrospinal fluid. The concentrations of these molecules are significantly elevated under pathological conditions. This is particularly evident in the serum of patients with liver diseases^3, 4^ and in the synovial and bronchoalveolar fluids of individuals with rheumatoid arthritis^5^ and idiopathic pulmonary fibrosis^6^, respectively. There is a strong correlation between the expression of ATX and several pathological conditions, such as inflammation^5^, obesity^7^, multiple sclerosis^8^, lung ^6^ and renal fibrosis^9^, liver steatosis^10^, cardiovascular disease^11^, bone erosion^12^, muscle regeneration^13^, as well as cancer growth and metastasis^14, 15, 16^. These association make ATX a compelling drug target. ATX is a multidomain molecule consisting of two SMB domains, a catalytic domain (phosphodiesterase domain, PDE), and a lasso loop connected to a nuclease domain (NUC)^17^. ATX adopts a compact structure, which results in the formation of distinct intramolecular compartments, including a tunnel and a hydrophobic pocket linked to the catalytic site^18^. The chemical research community is actively involved in developing more potent enzymatic inhibitors based on structural biology approaches ^19^. These inhibitors are currently classified into five categories based on their intramolecular location within ATX ^19, 20^.

LPA is a potent molecule with growth factor-like activities ^2^. However, it is also susceptible to catalytic degradation into monoacylglycerol by lipid-phosphate phosphatases expressed on the cell surface or by non-specific phosphatases present in biological fluids. The presence of these phosphatases serves as a biological defense mechanism against cell activation by LPA^21^. In this context, cargo transport proteins, such as albumin and gelsolin in the bloodstream, can uptake free LPA, thereby preserving its structural integrity^22, 23^. Consequently, ATX is now considered as a chaperone for LPA as well ^24^. Additionally, the demonstration of ATX’s interaction with cell-surface adhesion molecules has led to the hypothesis that local production of LPA near its receptors may offer an effective mode of cell activation ^25^. However, this hypothesis still requires direct validation at the pathophysiological level.

All five isotypes of ATX (α, β, γ, δ, and ε) contain Somatomedin B1/2 (SMB1/2) domains, including the essential amino acids E110 and H119, which are involved in the binding of ATX to β3-integrins ^18^. Nevertheless, the specific sequence of β3-integrins that interacts with ATX remains to be identified. ATXα binds to heparan sulfate chains via a 52-amino-acid polybasic insertion sequence ^26^, although the functional consequences of this interaction have not yet been determined. The binding of ATXβ to the syndecan-4 core protein has been demonstrated in previous studies, which have also shown that this interaction influences cell proliferation and cancer cell metastasis ^27^. However, the exact sequence on syndecan-4 that interacts with ATXβ has not been characterized. Furthermore, ATXβ has been shown to interact with α4β1 integrin, thereby regulating the migration of CD4+ T lymphocytes into secondary lymphoid organs. This evidence supports the physiological function of ATX binding to the surface of T lymphocytes ^28^. Nevertheless, the precise domains involved in the interaction between ATX and β1-integrin have yet to be identified.

The present study identified the ATX-fractalkine-like-domain (ATX-FLD) and the β1-integrin-fractalkine-binding-site (β1-IFBS) as the protein motifs involved in the interaction between ATX and β1-integrin. Additionally, our study demonstrated that LPA anchoring influences the three-dimensional structure of ATX, thereby exposing the ATX-FLD and enabling the binding of specific antibodies. A distinctive feature of these antibodies is that, although they do not interfere the lysoPLD activity, they inhibit the specific binding of ATX to cells expressing β1-integrins. Furthermore, the results from in vivo experiments suggest that targeting the interaction domains between ATX and β1-integrin may offer a novel therapeutic strategy for the treatment of inflammation and osteoarticular degradation.

## RESULTS

We previously demonstrated that cancer cells expressing high levels of α_V_β_3_ integrin exhibit a higher propensity to bind to ATX compared to cells expressing low levels. These results indicated that β_3_-integrin is involved in the binding of ATX to the cell surface ^29^. Expanding our studies to other cancer cell lines, we found that human osteosarcoma KHOS cells are a remarkable exception ^27^. Despite exhibiting a dramatically low level of α_V_β_3_ integrin expression, these cells showed the highest binding capacity for ATX compared to all other cell lines previously tested ^27^. Therefore, we conducted further studies to identify the additional factors that dominate the interaction of KHOS cells with ATX. It was demonstrated previously that the binding of ATX to non-adherent cells, specifically CD4+ T lymphocytes, is mediated by β1-integrin^28^. Consequently, we performed competition binding assays of KHOS cells on ATX using a series of anti-human integrin blocking antibodies (Figure 1A). In contrast to the anti-β2 and anti-αVβ5 integrin antibodies, which had no effect, and the anti-αVβ3 integrin antibody, which showed minimal inhibition ^27^, the anti-β1 integrin blocking antibody exhibited a significantly pronounced inhibitory effect, reducing KHOS cell binding to ATX by 97% (Figure 1A). To further confirm the general involvement of β1-integrin in the binding of adherent cells to ATX, we utilized calvaria-derived mouse osteoblasts (COB), where the β1-integrin gene had been knocked out (COB β1^−/-^), as well as β1-integrin-expressing parental cells (COB β1^fl/fl^) ^30^ in an ATX adhesion assay. In contrast to COB β1^fl/fl^ cells, which demonstrated a high binding capacity to ATX, COB β1^−/-^ cells showed a significantly reduced binding capacity (Figure 1B). These findings suggest that β1-integrin-mediated cell adhesion to ATX may represent a universal mechanism shared by both non-adherent and adherent cells. Moreover, these results position KHOS cells as a distinctive model for investigating the molecular mechanisms underlying the ATX interaction with β1-integrin, given the minimal involvement of β3-integrin in this process.

**Figure 1:**
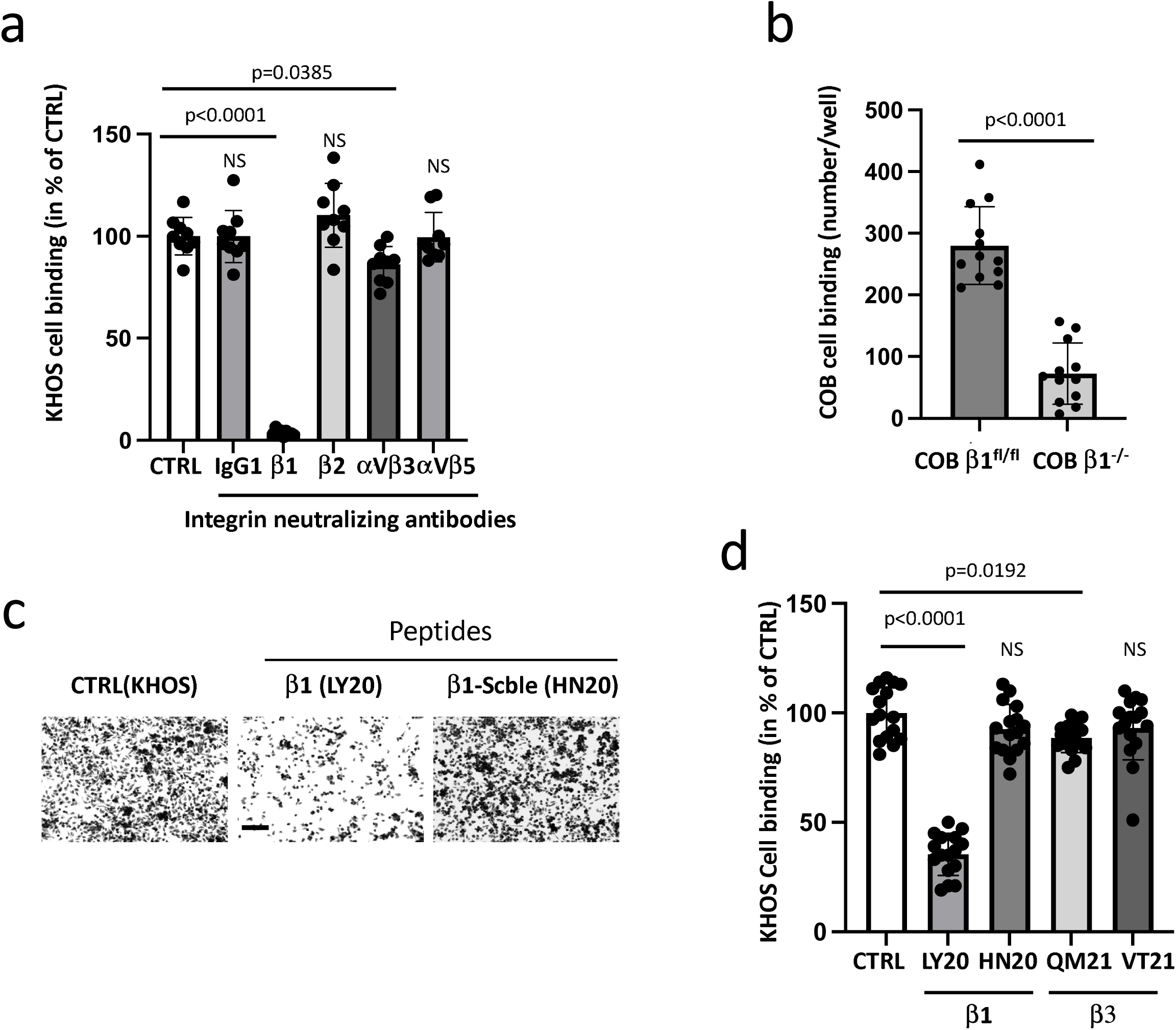
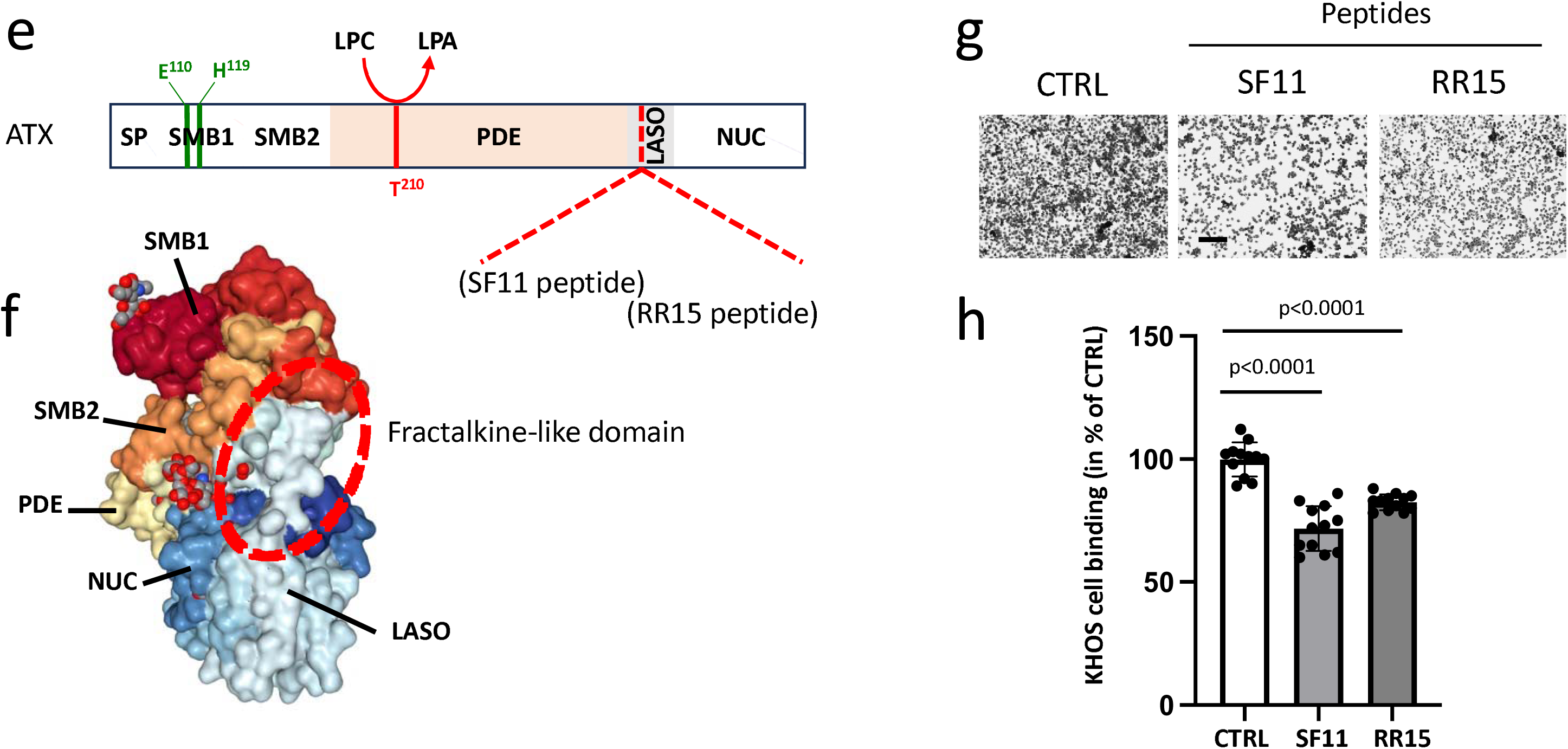
β1-integrin-fractalkine-binding site mediates cell binding to ATX via the ATX-fractalkine-like domain. **(A, C, D, G, H)** KHOS cells were preincubated for 1 h in the presence of Tyrode/HEPES buffer (CTRL), or **(A**) isotypic antibody (IgG1) or neutralizing antibody anti-β1 integrin, anti-β2 integrin, anti-α_V_β_3_ integrin or anti-α_V_β_5_ integrin (10 μg/mL) or **(C,D)** β1-LY20, β1-HN20, **(D)** β3-QM21, β3-VT2, **(G, H)** SF11 or RR15 peptides (50µM) before seeding for 1 hour on ATX (70µg) coated plates. **(B)** COB β1^fl/fl^ and COB β1^−/-^ were seeded for 1 hour on ATX (70µg) coated plates. The data are presented as the mean ± s.d of cell per well. **(C, G)** Representative images of KHOS cell adhesion plates. Scale bar represents 200 μm. **(E)** Primary structure of ATX depicting specific domains (SP: signal peptide, SMB1: somatomedin-like domain 1, SMB2: somatomedin-like domain 2, PDE: phosphodiesterase domain, LASO: laso domain, NUC: nuclease domain). Amino acid E110 and H119 are involved in β3-integrin binding. Amino acid T210 is necessary for lysoPLD activity leading to LPA synthesis from LPC. Sequences and locations of SF11 and RR15 peptides are indicated in ATX primary structure. **(F)** The fractalkine-like domain is highlighted in red dotted line on ATX three-dimensional structure (http://www.rcsb.org/3d-view/3NKP). **(A, B, D, E)** The results were obtained from three to four biologically independent experiments with n=3 to 4 replicates. NS: not significant. **(A, D, H)** The data are presented as the mean ± s.d of cells in % of CTRL and statistical analysis by ordinary one-way with Dunnett’s multiple comparisons test. **(B)** The data are presented as the mean ± s.d of cell number/well and statistical analysis by Mann-Whitney test. Source data are provided as a Source data file.

The primary sequence of ATX contains the canonical RGD motif, which is known to mediate binding to β3-integrin ^31^. However, both our studies and those of others have demonstrated that cell binding to ATX occurs independently of the RGD sequence ^18, 29^. Therefore, we sought to identify sequence homologies between ATX and RGD-independent ligands of β1-and/or β3-integrins. Among the identified candidates, fractalkine was shown to bind to the amino acid sequence 275-294 of β1-integrin and 267-287 of β3-integrin ^32^. To investigated this interaction, we performed a competition binding assay with ATX using KHOS cells, in the presence of synthetic peptides corresponding to the β1-integrin-fractalkine binding site (β1-IFBS, peptide LY20) and the β3-integrin-fractalkine binding site (β3-IFBS, QM21), as well as corresponding scrambled peptides (β1-HN20 and β3-VT2) (Figure 1C, D). Consistent with the minimal involvement of β3-integrin in KHOS cell adhesion to ATX (^27^ and Figure 1A), the β3-QM21 peptide showed a low but still significantly 11% inhibition of KHOS cell adhesion. In contrast, the β1-LY20 peptide, but not the scrambled β1-HN20 peptide, led to a significant 65% inhibition of KHOS cell binding to ATX (Figure 1C-D). These findings identify, for the first time, the amino acid sequences 275-294 of β1 integrin and 267-287 of β3 integrin, as interaction sites for ATX.

As a complementary approach, we searched for sequence homologies between ATX and fractalkine. We identified the sequence SLNHLLRTNTF on ATX that shares 63.6% identity over 11 amino acids with fractalkine. To confirm the binding reciprocity, we conducted additional binding assays with ATX using synthetic peptides (SF11 and RR15), which overlap the fractalkine-like domain (Figure 1E) located within the LASO domain of ATX (Figure 1E, F). The SF11 and RR15 peptides both exhibited a significant inhibitory effect on KHOS cell binding to ATX, with inhibition rates of 28% and 18%, respectively (Figure 2G, H). In conclusion, these results demonstrate that the interaction of the ATX-fractalkine-like domain (ATX-FLD) and the β1-integrin-fractalkine-binding site (β1-IFBS) plays a critical role in KHOS cell binding to ATX.

**Figure 2:**
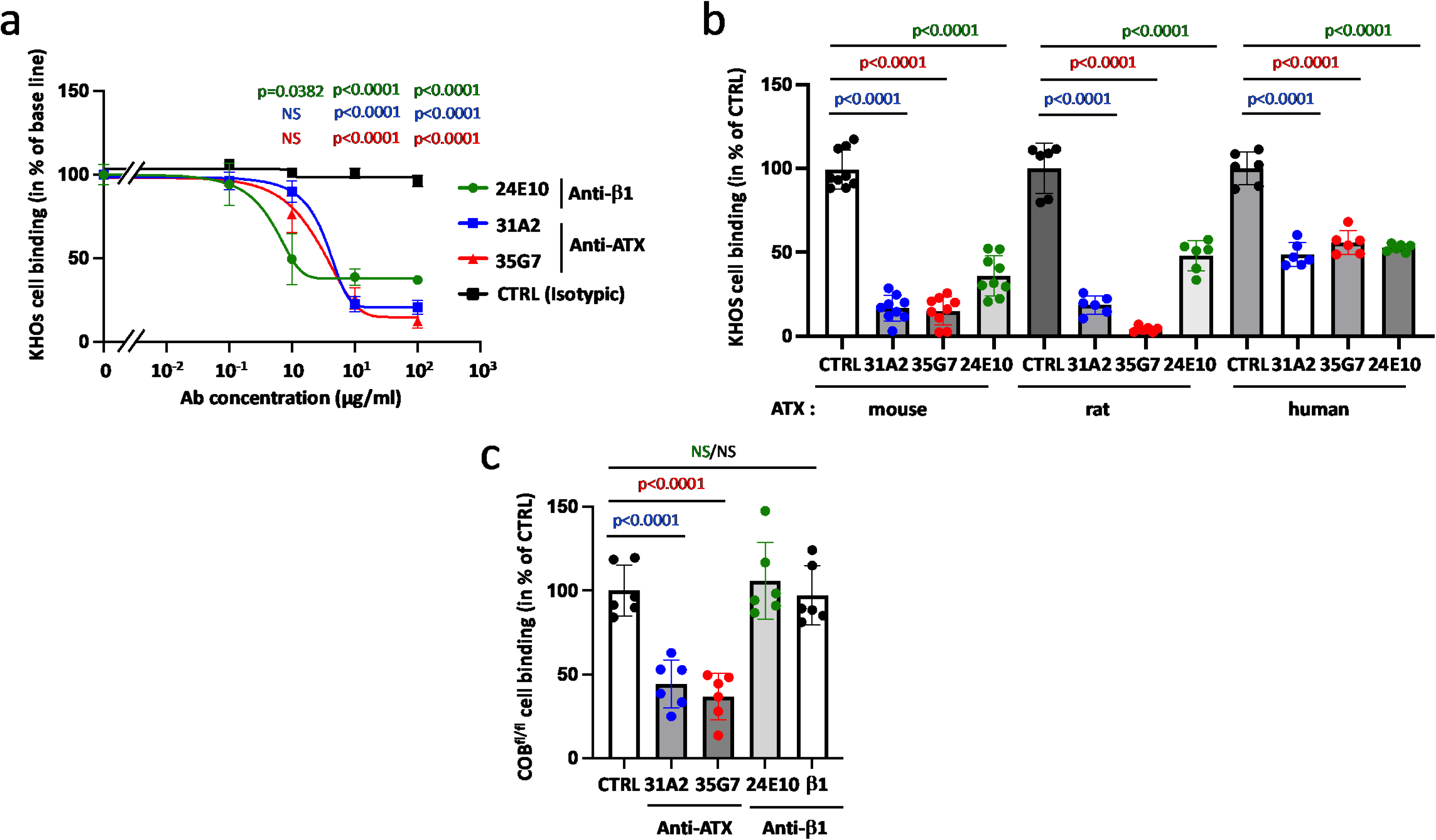
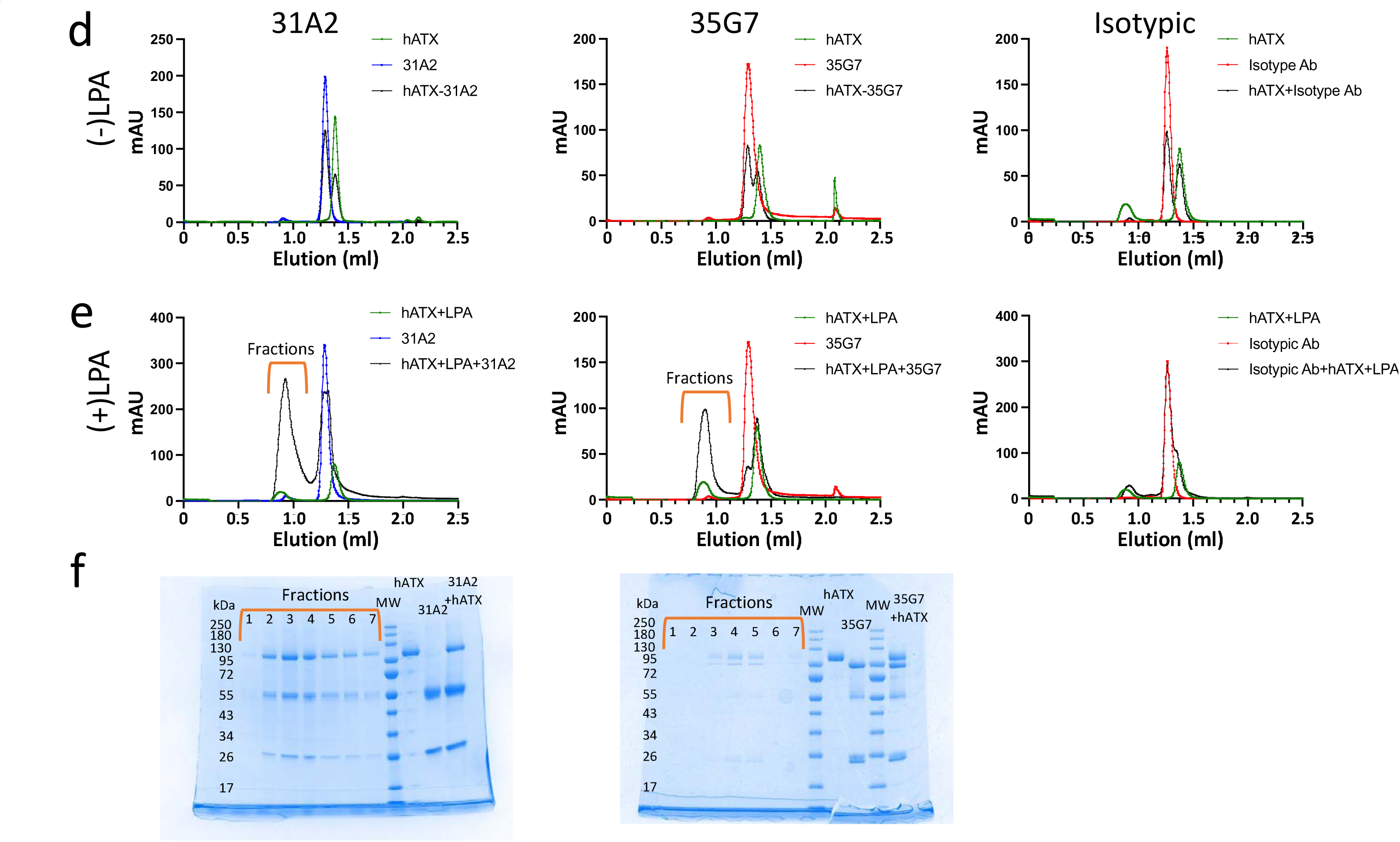
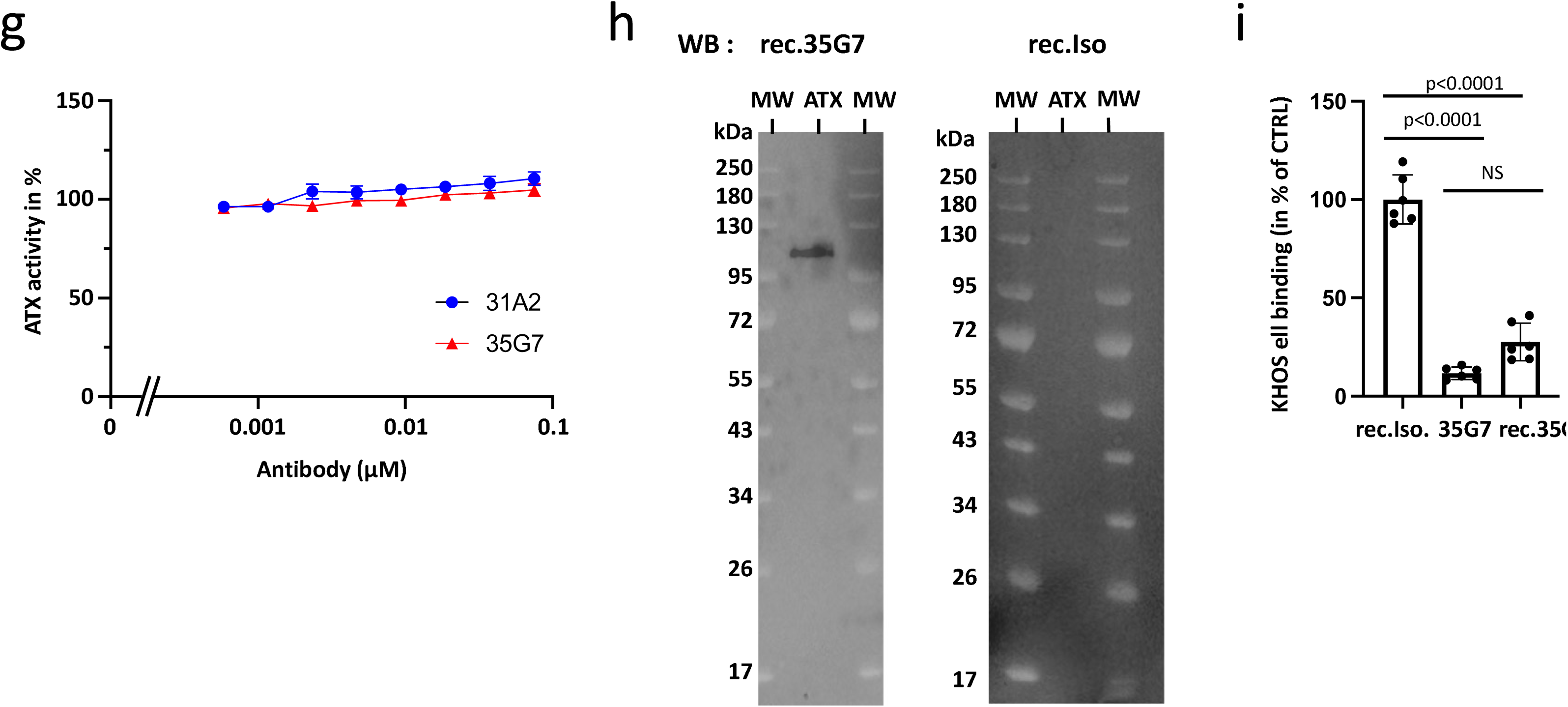
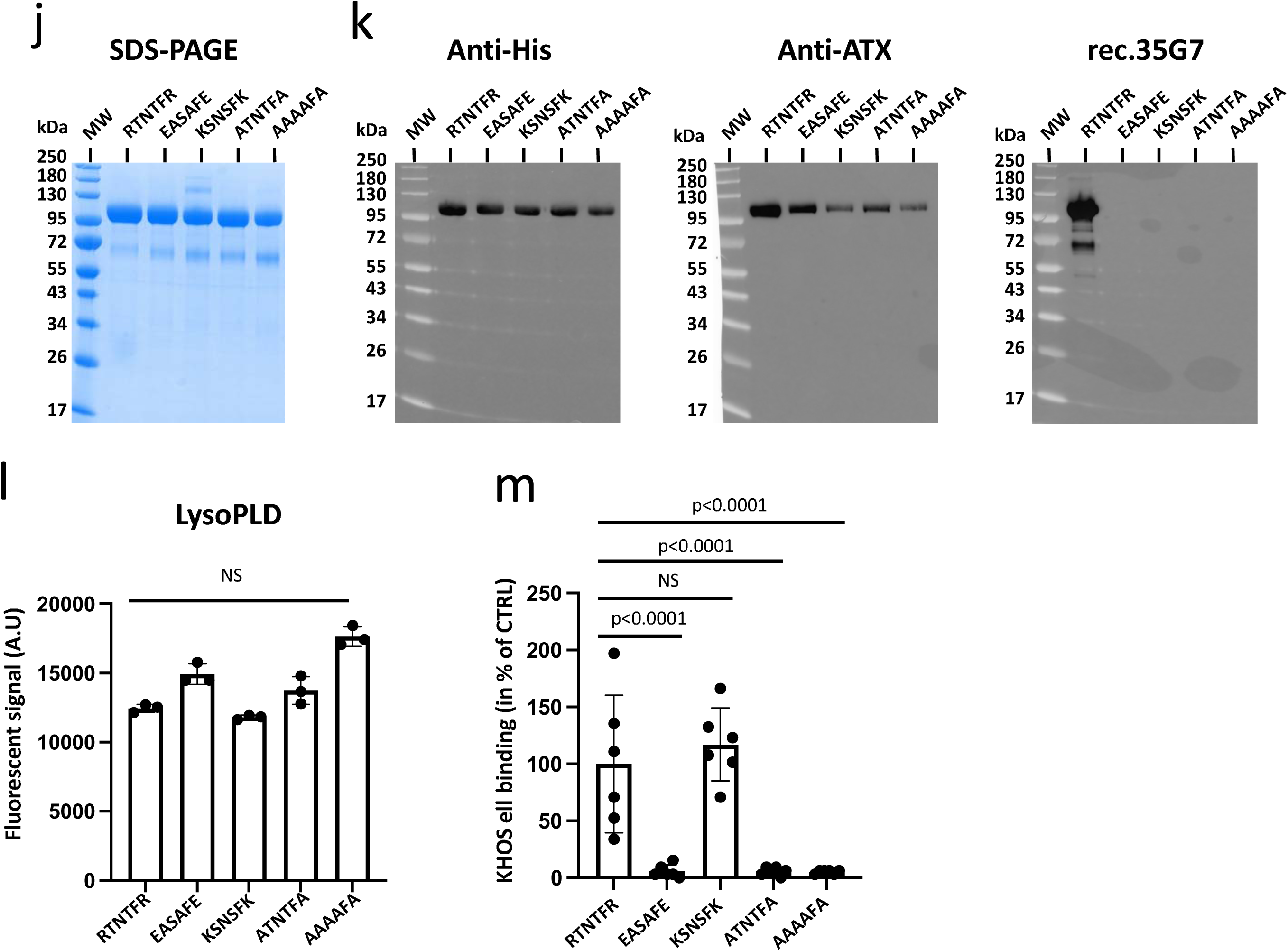
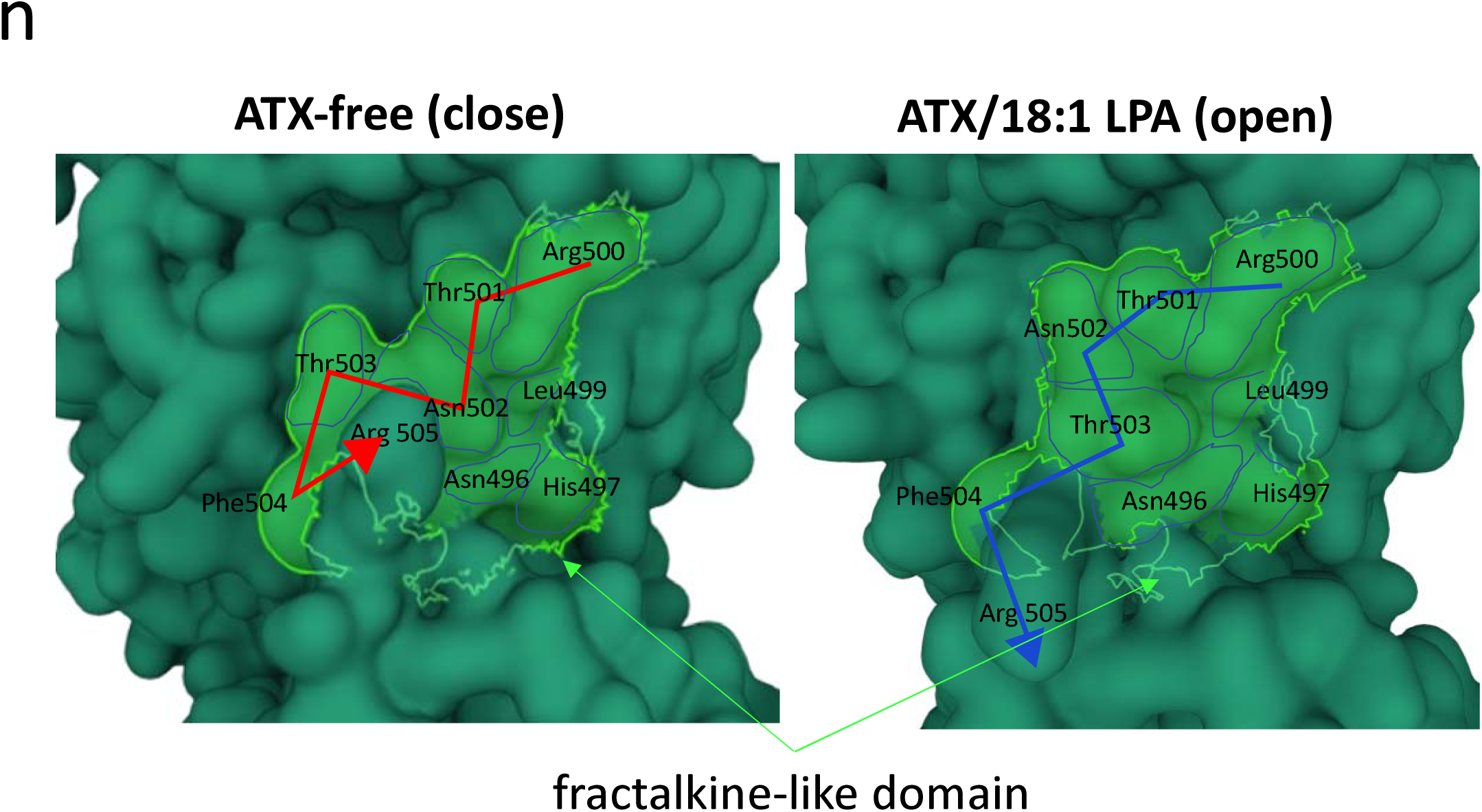
Characterization of anti-β1-integrin-fractalkine-binding-site (anti-β1.IFBS) and anti-ATX-fractalkine-like-domain (anti-ATX-FLD) monoclonal antibody inhibitory actions on cell binding to ATX. **(A)** Concentration-response curves of antibodies on KHOS cell binding to mouse ATX. KHOS cells were preincubated for 1 h in the presence of indicated concentrations of Isotypic (CTRL) or anti-β1-IFBS (24E10) or anti-ATX-FLD (31A2, 35G7) antibodies and seeded on plates coated with mouse ATX (70ng). **(B)** Inhibitory action of anti-β1.IFBS and anti-ATX-FLD antibodies on KHOS cell adhesion on mouse, rat or human ATX. KHOS cells were preincubated with for 1 h in the presence of Isotypic (CTRL), 24E10, 31A2 or 35G7 (10µg/ml) antibodies and seeded on plates coated with mouse, rat or human ATX (70ng). **(C)** Effect of anti-human β1, anti-β1-IFBS and anti-ATX-FLD antibodies on COB^fl/fl^ cell adhesion on ATX. COB β1^fl/fl^ cells were preincubated for 1 h in the presence of Isotypic (CTRL), 24E10, 31A2, 35G7 or anti-human CD29 (β1) antibodies (10µg/ml) and seeded on plates coated with mouse ATX (70ng). **(A)** The data are presented as the mean ± s.e.m of cells in % of base-line (absence antibody). The results were shown from three biologically independent experiments with n=3 replicates. Statistical analysis by two-way ANOVA with Turkey’s multiple comparisons test. **(B, C)** The data are presented as the mean ± s.d of cells in % of CRTL. The results were shown from two to three biologically independent experiments with n = 3 replicates. Statistical analysis by ordinary one-way ANOVA with Dunnett’s multiple comparisons test. **(D-F)** LPA controls 31A2 and 35G7 binding to ATX. Size Exclusion Chromatography of antibodies, human ATX and complex mixture. Elution was done in Tris 50mM, NaCl 150 mM pH 7.4 using a Superdex 3.2 / 300 increase 200 on a microAKTA purifier. Antibody 31A2, 35G7 or isotypic were in 1.2 ratio with antigen and were incubated 1h at 4 degrees **(D)** in absence (-LPA) or **(E-F)** in presence (+LPA) of 18:1 LPA before injection. **(F)** Coomassie blue SDS page gel in reducing conditions. Eluted peaks were loaded on position 1 to 7. **(G)** Anti-ATX-FLD antibodies do not affect ATX LysoPLD activity. LysoPLD ATX activity was measured as released choline by 50 μM 18:1 LPC hydrolysis in the presence of increasing concentration of antibodies 31A2 and 35G7. ATX concentration in the assay is 30 nM. Errors bars represent s.e.m from biological triplicate. **(H)** Western blotting detection of rat ATX (100ng) in reducing condition by rec.35G7 but not by rec.Iso. **(I)** Anti-ATX-FLD rec.35G7 antibody inhibits KHOS cell binding to ATX. Conditions and analyses are identical as described in (B). **(J, K)** Mutations in ATX-FLD affect rec.35G7 antibody recognition but not LysoPLD activity**. (J)** SDS page and **(K)** western blot analysis of rat ATX mutants. **(J)** Recombinant ATX (2 μg) were loaded on SDS page and stained using instant blue. **(K)** Western blotting detection of ATX (100ng) in reducing condition by antibody ab18184 (anti-His), MABT1350 (anti-ATX) but not by anti-ATX-FLD (rec.35G7). **(L)** LysoPLD ATX activity measured as released choline by 50 μM 18:1 LPC and recombinant ATX mutants (30 nM). Errors bars represent s.e.m from n=3 biological replicates. **(M)** KHOS cells were seeded on plates coated with recombinant wild-type (RTNTFR) or mutants (EASAFE, KSNSFK, ATNTFA, AAAAFA) of rat ATX (200ng). **(N)** The presence of 18:1 LPA induces structural modification of the fractalkine-like domain of ATX. 3D views from the RCSB Protein Data Bank free access database (https://www.rcsb.org) of murine ATX in the absence of any compound (ref. 2XR9, ATX-free) showing a “closed” conformation of the fractalkine-like domain, whereas co-crystallised murine ATX with 18:1 LPA (ref. 3NKP, ATX/18:1 LPA) shows an “open” conformation of this domain. Source data are provided as a Source data file.

Subsequently, we generated a series of monoclonal antibodies by immunizing mice with synthetic peptides, leading to the selection of one anti-β1-IFBS antibody (24E10) and two anti-ATX-FLD antibodies (31A2 and 35G7). Each antibody demonstrated a significantly concentration-dependent inhibition of KHOS cell adhesion to mouse ATX (Figure 2A). The sequence of ATX is highly conserved across species. However, since all prior ATX binding experiments were performed using mouse ATX, we sought to determine whether ATX from different species might affect the blocking capacities of these monoclonal antibodies. First, the anti-β1-IFBS 24E10 antibody exhibited comparable inhibitory efficacy against KHOS cell adhesion to mouse (64%), rat (52%) or human (48%) ATX (Figure 2B). The anti-ATX-FLD antibodies, 31A2 and 35G7, demonstrated greater potency in inhibiting KHOS cell adhesion to murine ATX compared to human ATX, with 31A2 showing 85% inhibition on both mouse and rat ATX, and 35G7 exhibiting 95% and 85% inhibition on rat and mouse ATX, respectively (Figure 2B). Although the complete expression pattern of β-integrins in COB cell remains unknown, COB β1^fl/fl^ cells demonstrated suitability for ATX binding assays (see Figure 1B). These cells were subsequently used to evaluate the inhibitory potential of the monoclonal antibodies on mouse cells. We observed that the 31A2 and 35G7 antibodies also inhibited the binding of COB β1^fl/fl^ cells to mouse ATX (Figure 2C), although the inhibition (56%–64%) was less than that observed in human KHOS cells (Figure 2B). In stark contrast, 24E10 antibody did not inhibit COB β1^fl/fl^ cell adhesion to mouse ATX, a result that was similarly observed with the specific anti-human β1-integrin antibody (CD29), suggesting that action of the 24E10 antibody is restricted to human β1-expressing cells.

Subsequently, we investigated the direct interaction between the 31A2 and 35G7 antibodies and soluble human ATX (hATX) using a size exclusion chromatography assay. Unexpectedly, mixing of hATX with either 31A2 or 35G7 antibody in solution resulted in the recovery of the individual proteins in separate elution fractions, indicating the absence of complex formation between ATX and each antibody (Figure 2D). In stark contrast, preincubation of ATX with LPA prior to mixing with either antibody led to the appearance of an early elution fraction peak (Figure 2E), which contained both ATX and the corresponding antibody (Figure 4F). The absence of this early elution peak when ATX-LPA was mixed with the isotypic control antibody further supports the binding specificity of 31A2 and 35G7 antibodies for soluble ATX only after priming with LPA (Figure 2E). This result suggests that the binding of the 31A2 and 35G7 antibodies to soluble ATX is dependent on the exposure of a cryptic epitope induced in the presence of LPA. Furthermore, the incubation of either 31A2 or 35G7 antibodies with soluble ATX and LPC did not interfere with lysoPLD activity (Figure 2G). These findings indicate that 31A2 and 35G7 antibodies specifically affect the binding of ATX to the cell surface via the β1-integrin, without affecting ATX’s function in producing LPA. We then generated a recombinant form of antibody 35G7 by transferring the variable sequence on an Fc-silent backbone (rec.35G7) for potential therapeutic use in preclinical mouse models. First, using western blotting, we confirmed that, unlike the recombinant isotype antibody (rec.Iso), rec.35G7 was capable of recognizing the denatured and reduced form of ATX (Figure 2H). Subsequently, we demonstrated that rec.35G7 exhibited the same inhibitory effect as the native 35G7 monoclonal antibody on KHOS cell adhesion on ATX (Figure 2I). Moreover, we generated a series of four recombinant forms of the rat-ATX containing distinct mutations in the fractalkine-like domain (RTNTFR) (Figure 2J). The mutants were designed to either preserve the electrostatic charge of the amino acid sequence (KSNSFK), exhibit an entirely opposing charge (EASAFE), harboring a single R-to-A substitution (ATNTFA) or display an almost complete A-substituted domain (AAAAFA). As the recombinant proteins were fused to a 6His-tag, enabling affinity purification via nickel column chromatography, all wild-type and mutant proteins were detected by western blotting using anti-His antibodies (Figure 2K). Furthermore, all proteins were recognized by the anti-ATX monoclonal antibody MABT1350 (Figure 2K). In stark contrast, rec.35G7 exhibited specificity for the wild-type ATX (Figure 2K) but failed to detect any of the mutants in western blotting. Additionally, all mutants retained intact lysoPLD activity (Figure 2L) indicating that the integrity of the RTNTFR domain is dispensable for ATX lysoPLD activity. Notably, all these mutants - excepted KSNSFK - were unable to support KHOS cell adhesion (Figure 2M) suggesting that, beyond the sequence itself, the proper electronic charge of the RTNTFR domain is crucial for ATX cell adhesion.

The three-dimensional structure of ATX was elucidated over a decade ago (Jens et al., 2011). Since then, numerous structural biology studies have been conducted to characterize the intramolecular localization of lysoPLD inhibitors ^19^. To identify potential structural changes of ATX due to the presence of LPA, we first compared the only two murine structures of ATX free of any compound and ATX co-crystallized in the presence of 18:1 LPA, as deposited in the open-access RCSB Protein Data Bank (https://ww.rcsb.org) (Figure 2N). This comparison revealed a notable conformational alteration in the fractalkine-like domain of ATX, which adopted a closed-type three-dimensional (3D) conformation in the absence of any compound and an open-type 3D conformation in the presence of 18:1 LPA. These findings support the hypothesis that the 35G7 epitope exists in a cryptic form in the compound-free state that become unmasked in the presence of 18:1 LPA.

Next, we assessed the effect of rec.35G7 in mice with collagen-induced arthritis (CIA). Mice received intraperitoneal injections of rec.35G7 and rec.Iso antibodies (1 mg/kg) twice a week from day 20 (the day of collagen immunization boost) until day 30 (Figure 3A). The administration of rec.35G7 did not result in any noticeable changes in body weight, a finding consistent with the observations in the vehicle and rec.Iso antibody-treated groups (Figure 3B). In stark contrast, animals treated with rec.35G7 exhibited a significant reduction in clinical arthritis scores in comparison to those receiving vehicle and rec.Iso antibodies (Figure 3C). Furthermore, rec.35G7 treatment led to a decreased incidence of severe knee erosion (28.5%) compared to vehicle-treated (71.5%) and rec.Iso-treated animals (71.5%), as assessed by microCT analysis at both knee and ankle sites (Figure 3). These findings indicate that rec.35G7 treatment may offer protection against osteoarticular inflammation and bone degradation in the context of rheumatoid arthritis. To further evaluate the clinical potential of rec.35G7 in the rheumatoid arthritis, we conducted a side-by-side comparison of its effects with those of the current standard-of-care anti-TNFα antibody, infliximab, in the CIA model. As previously found, rec.35G7 did not induce any adverse effect on animal body weight, a finding similarly noted for infliximab treatment (Figure 3F). Notably, both rec.35G7 and Infliximab resulted in a comparable reduction in arthritis scores (Figure 3G). These findings suggest that rec.35G7 may represent a promising novel immunotherapy for chronic inflammatory rheumatisms.

**Figure 3:**
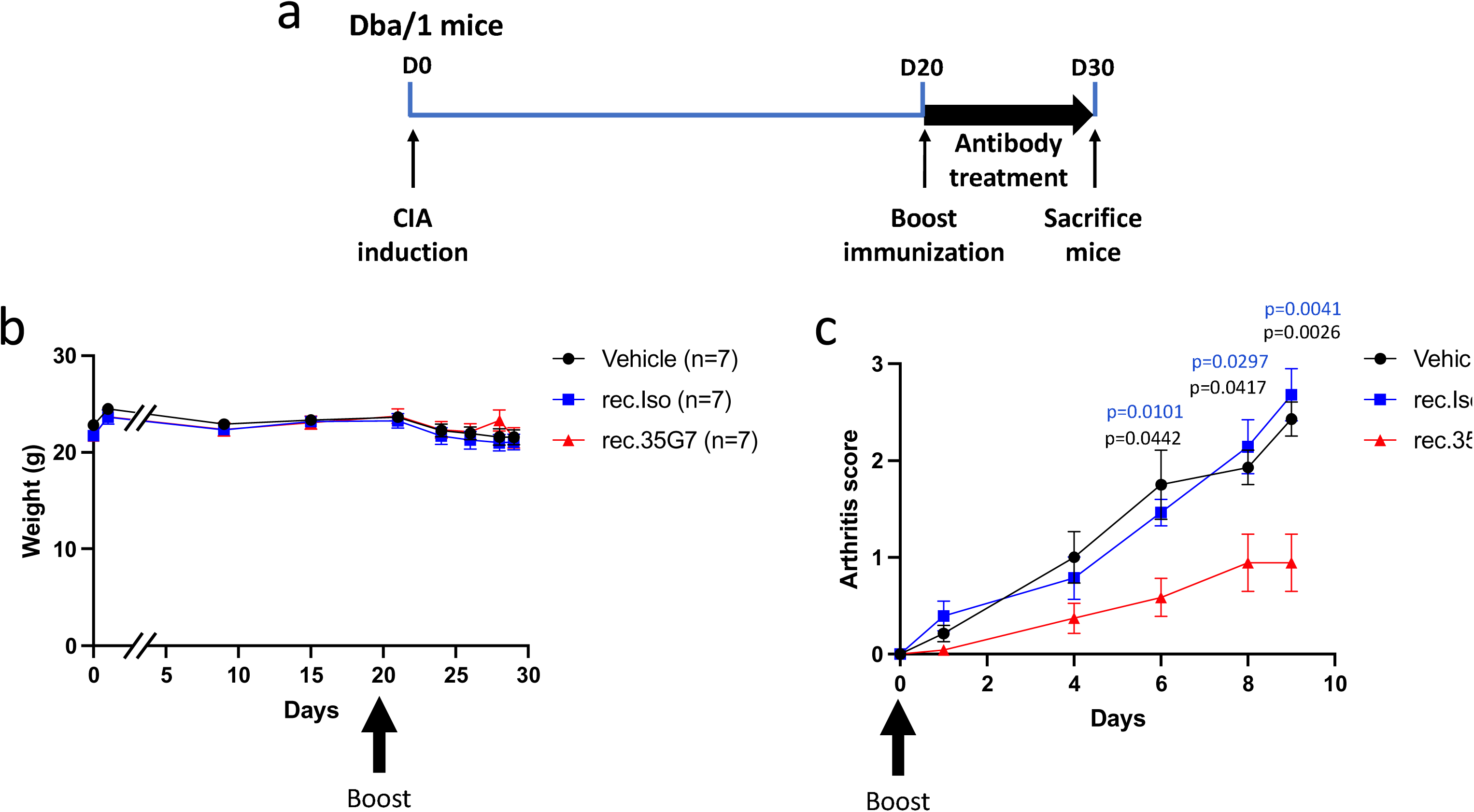
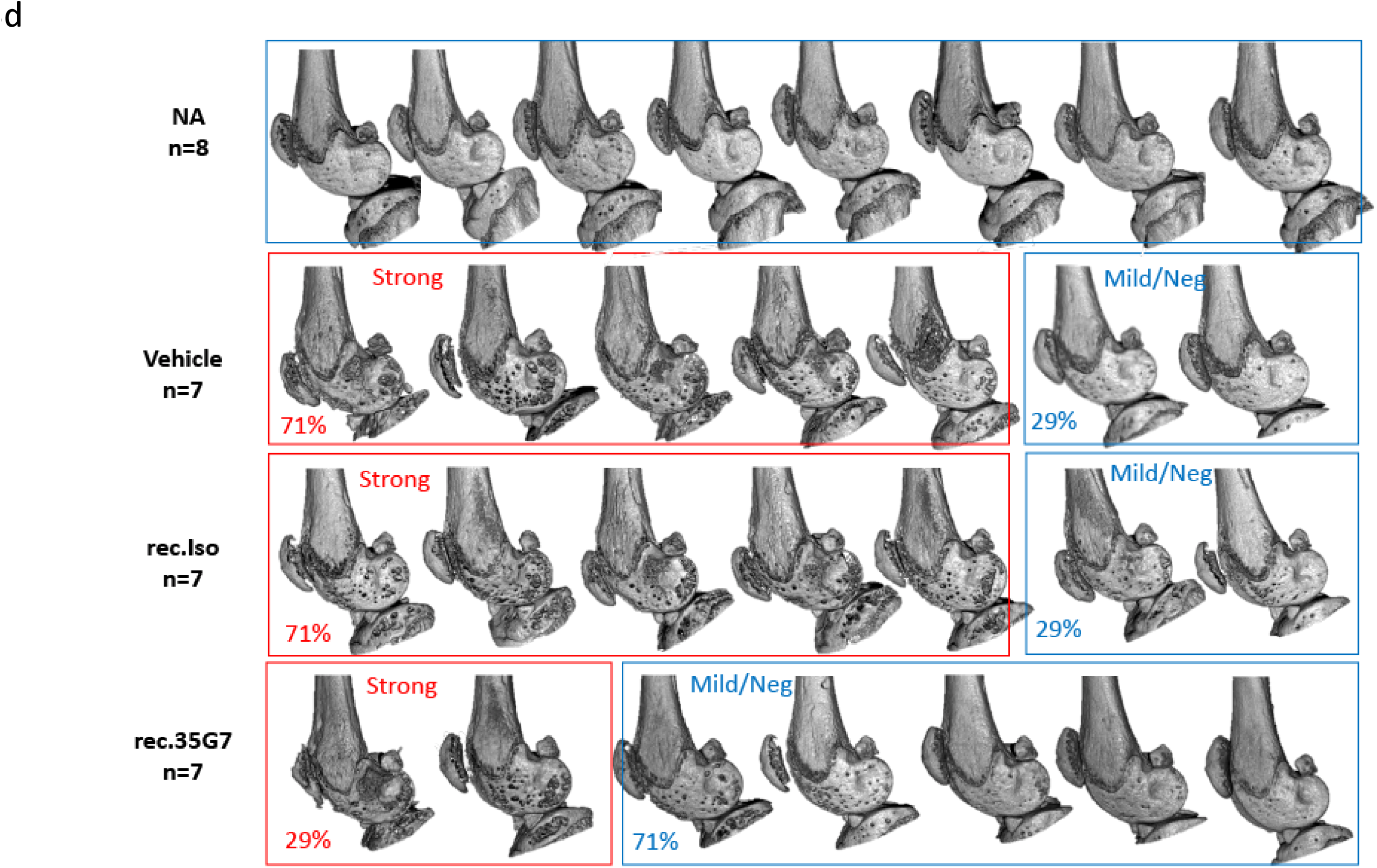
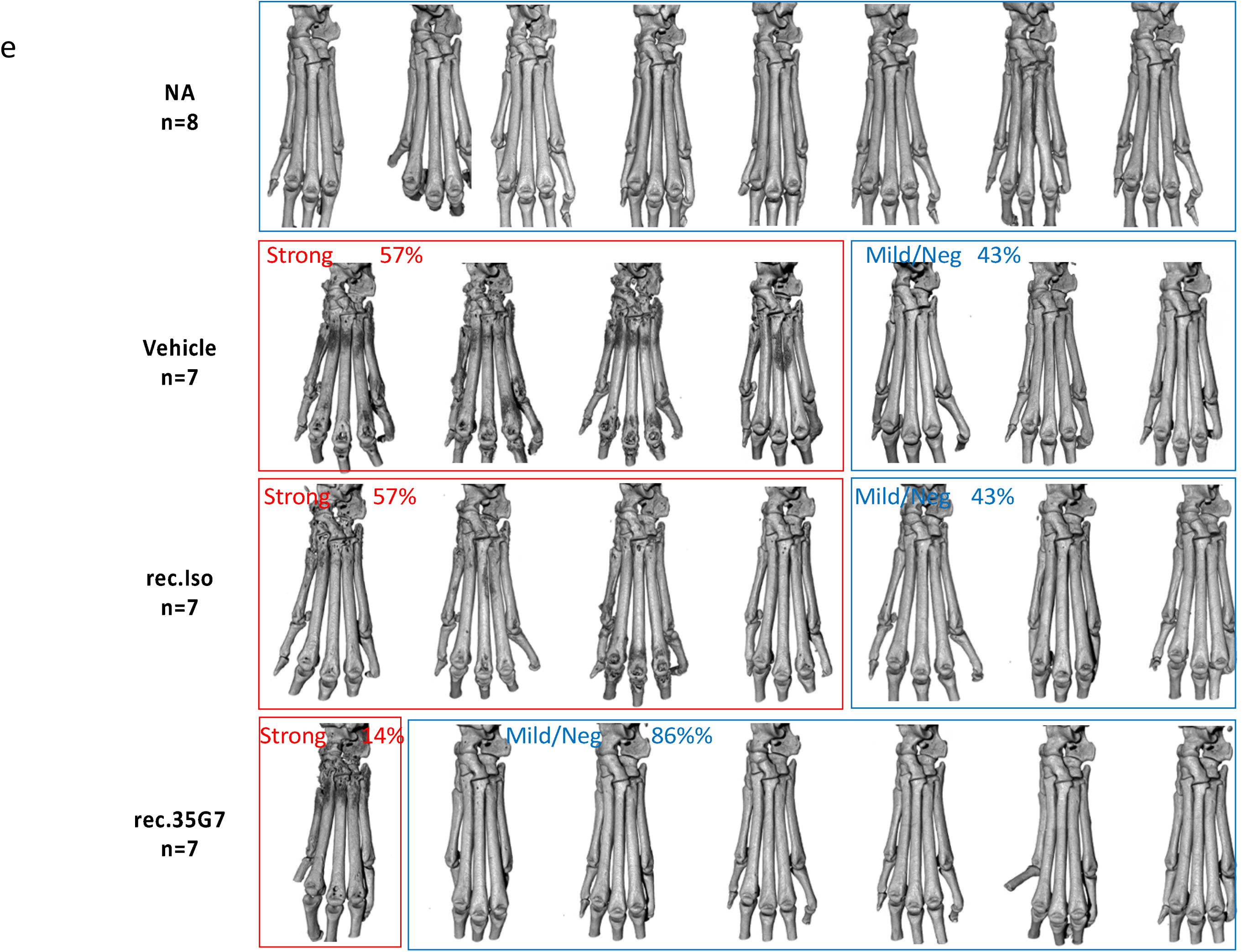
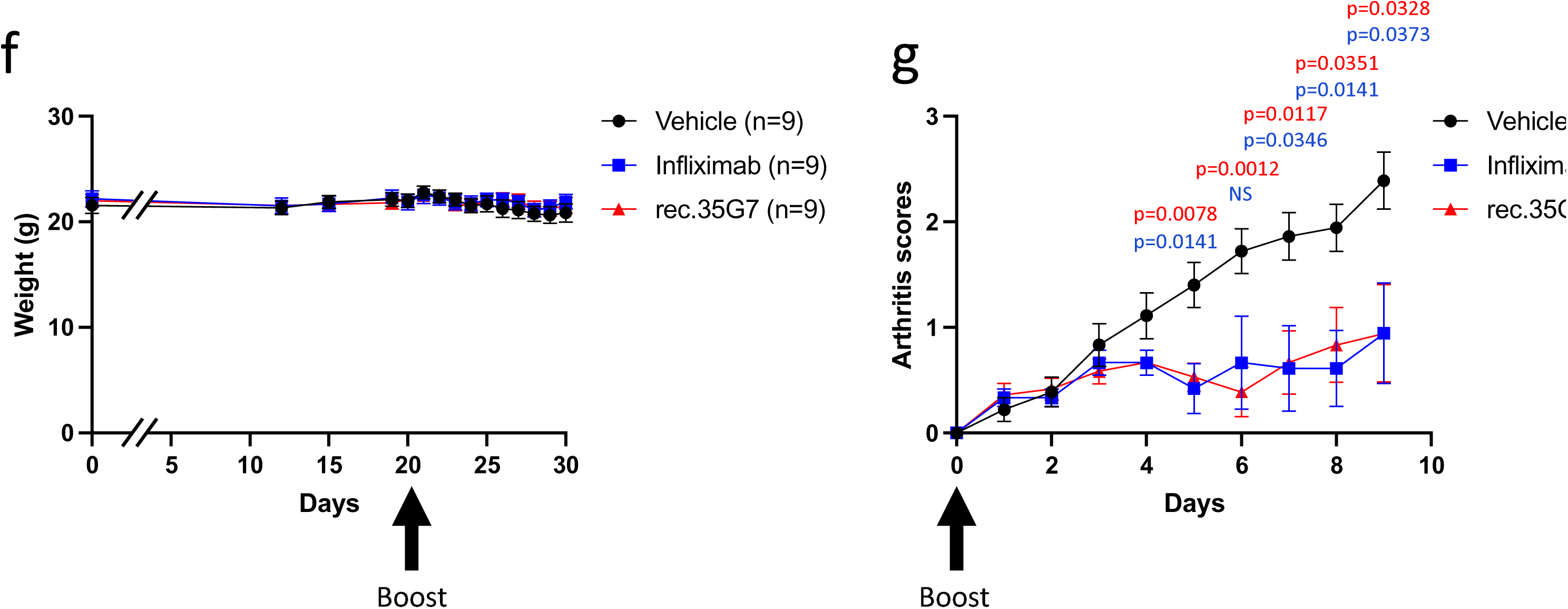
Treatment with rec.35G7 antibody attenuates collagen induced arthritis. **(A)** Schematic representation of immunization and antibodies administration schemes. DBA/1 mice subjected to CIA were treated intraperitoneally with vehicle, rec.Iso (1 mg/kg), or rec.35G7 (1 mg/kg) twice a week from day 20 until day 30. **(B)** Time dependent animal weight after primary immunization with type II collagen. **(C)** Time dependent arthritis scores after boost immunization with type II collagen. **(B,C)** Errors bars represent s.e.m from n=7 animals per group. Representative microCT images of **(D)** knees and **(E)** ankles (front view: upper panels; side view : lower panels) of non-arthritic animals (NA), and Vehicle, rec.Iso and rec.35G7-treated mice. **(F-G)** DBA/1 mice subjected to CIA were treated intraperitoneally with vehicle, Infliximab (1 mg/kg), or rec.35G7 (1 mg/kg) twice a week from day 20 until day 30. **(F)** Time dependent animal weight follow-up after primary immunization with type II collagen. **(G)** Time dependent arthritis score follow-up after boost immunization with type II collagen. **(F,G)** Error bars represent s.e.m from n=9 animals per group. **(B,C,F,G)** Statistical analysis by two-way ANOVA with Turkey’s multiple comparisons test. Source data are provided as a Source data file.

**Figure 4:**
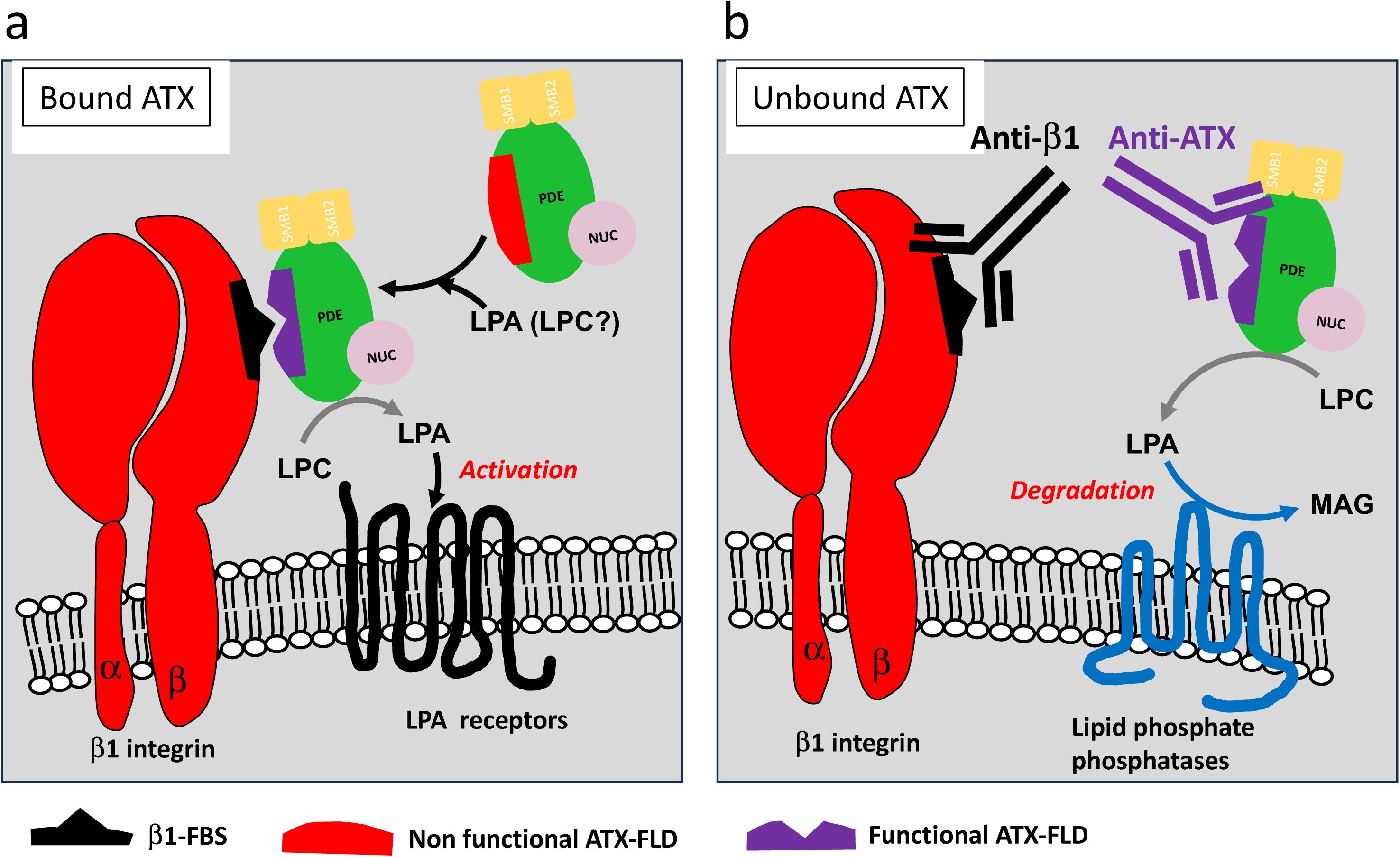
Novel anti ATX strategy interfering with the mechanism of ATX binding to β1-integrin. **(A)** The presence of LPA induces a conformation change in the fractlakine-like-domain (FLD) of ATX, from a non-functional (Red) to a functional (Purple) form allowing ATX to interact with the β1-integrin-fractalkine-bining site (β1-FBS in black), potentially favoring LPA receptor activation due to local LPA production. **(B)** Anti-β1-FBS (24E10) and anti-ATX-FLD (31A2, 35G7) antibodies interfere with ATX binding to β1-integrin. In this context, unbound ATX may accumulate in the extracellular compartment, moving the site of LPA production away from its receptors thereby promoting its degradation into monoacylglycerol (MAG) by lipid phosphate phosphatases.

## DISCUSSION

In this study, we identified a fractalkine-like domain on ATX (ATX-FLD) that mediates the binding of ATX to β1 integrin. Furthermore, we demonstrated that while monoclonal antibodies targeting ATX-FLD do not affect ATX lysoPLD activity, they alleviate inflammation and articular bone degradation in a CIA mouse model. Moreover, our study revealed that the recognition of soluble ATX by anti-ATX-FLD antibodies depends on the exposure of a cryptic epitope, which is unmasked following structural modifications of the Lasso domain in the presence of 18:1-LPA. ATX is a dual-function protein, acting both as an LPA-producing enzyme and as a chaperone in LPA signaling^33^. This function protects LPA from degradation by phospholipases and provides a privileged mechanism for LPA receptor activation. Our d-findings also indicate that the occupancy of the tunnel and hydrophobic channel by 18:1-LPA regulates ATX binding to β1-integrin. However, it remains to be determined whether similar structural modifications can be induced by other LPA species or by specific types of lysoPLD inhibitors occupying these intramolecular compartments.

Incubation of T lymphocytes with the lysoPLD-deficient ATX-T210A leads to a reduction in cell colonization of secondary lymphoid organs^28^. This is effects is attributed to a competitive binding with high endothelial venules-derived ATX to β1-integrin on T lymphocytes. The present study provides preclinical evidence that blocking the ATX pathway by targeting its binding to the cell surface at the site of chronic inflammation may have therapeutic value. Furthermore, our findings demonstrate that inhibiting the binding of ATX to the cell surface without affecting its lysoPLD activity represents an effective strategy for mitigating disease progression in a RA mouse model.

ATX is a secreted factor present in soluble form in biological fluids, including blood, where it plays a key role in regulating the LPA levels ^34^. However, the biological functions of both ATX and LPA at the systemic level remain unclear. The interaction of ATX with cell surface adhesion molecules, including β1 and β3 integrins, has been previously characterized ^18, 28, 29, 35^. This has led to the current hypothesis that the binding of ATX to adhesion molecules would favor the production of LPA in close proximity to its cell surface receptors, thereby facilitating cell activation^25^. Nevertheless, the specific domains involved in the interaction with β1 and β3 integrins remain to be identified. All ATX isoforms contain both a RGD and a LDV motif ^36^, which are recognized as canonical interaction sequences for β3 and β1 integrins, respectively^31, 37^. However, experimental evidence has demonstrated that these sequences do not mediate ATX binding to β3 and β1 integrins, respectively ^28, 29, 35^. Instead, mutations in the coding sequence for amino acids E110 and H119 have been shown to disrupt the binding of β3-integrin expressing cells^18^, suggesting that ATX interacts with β3-integrins through a non-canonical mechanism. By aligning ATX amino acid sequence with RGD-independent ligands of β1 and β3 integrins, we identified a fractalkine-like motif (SLNHLLRTNTF) within the Lasso domain of ATX. All mutations within the RTNTFR sequence of the rat ATX significantly impaired KHOS cell adhesion, excepted for the KSNSFK mutant. This finding suggests that, beyond the primary sequence, the proper electronic charge of the RTNTFR domain is essential for ATX-mediated cell adhesion. Furthermore, our study demonstrated that the fractalkine binding site on β1 and β3 integrins (IFBS) significantly contributes to ATX cell binding. The use of anti-ATX-FLD antibodies effectively inhibited human and mouse cell binding to both human and murine ATX. In contrast, the anti-β1-IFBS monoclonal antibody exhibited an inhibitory effect restricted to human cells, despite showing a similar level of inhibition on both human and murine ATX. Consequently, the therapeutic potential of this antibody could not be assessed in the present study, as it would require the use of either a humanized mouse model or anti-mouse β1-IFBS antibodies.

The tools generated in this study will provide valuable insights into the intricate molecular mechanisms governing ATX’s effects on immune cells, platelets, fibroblasts, cancer cells and others. ATX exhibits a repulsive action on eosinophil recruitment in pancreatic cancer ^38^, as well as on CD8+ T lymphocytes in melanoma^39^. In contrast, it facilitates the migration of Jurkat T cells and the invasion of CD4+ T lymphocytes into secondary lymphoid organs^40^. Furthermore, ATX has been shown to promote inflammation in conditions such as lypopolysaccharide-induced sepsis^41^, idiopathic pulmonary fibrosis (IPF) ^6^ and rheumatoid arthritis (RA)^5^. The complex mode of action of ATX may depend on the specific types of LPA receptors involved in these processes, as evidenced by the prevalence of downstream signaling activation of LPA5 and LPA6 in melanoma ^39, 42^ and of LPA1 in IPF ^43^, RA^44^ and vasculitis^45^. Additionally, recent studies have demonstrated that ATX displays functional selectivity for P2Y-type LPA6 over EDG-type LPA1^33^. Therefore, in addition to its dual protein function, the biological activity of ATX relies on a multistep signaling cascade linked to its ability to produce LPA, which drives cell-specific activation via a distinct set of THE LIGAND receptor subtypes. Our findings suggest that particular subclasses of β-integrins involved in ATX binding may also contribute to this complexity. Analogous to the formation of a ternary complex for fractalkine signaling with CX3CR1 ^32^ and uPAR signaling with vitronectin^46^, it can be hypothesized that the specificity of the β-integrin creates a platform that connects ATX-derived LPA to the activation of specific subtypes of LPA receptors.

In conclusion, the results of this study provide a more comprehensive understanding of the molecular mechanisms that regulate ATX biological effects. In the context of RA, this is further supported by an LPA-dependent interaction of ATX with β1-integrin expressing cells. These findings also pave the way for the development of novel therapeutic strategies targeting ATX, which can be tailored to specific functions of ATX while maintaining its lysoPLD activity. This approach would circumvent potential off-target effects associated with the systemic inhibition of lysoPLD activity.

## MATERIALS AND METHODS

### Cell culture and reagent

Human osteosarcoma cell lines KHOS were obtained from the American Type Culture Collection (ATCC; Gaithersburg, MD, USA). Immortalized primary mouse calvaria osteoblasts expressing of β1 integrin (COB-β1^fl/fl^) and deleted for the gene encoding β1 integrin (COB-β1^−/-^) were a generous gift from Dr. D. Bouvard ^30^. All cells were cultured in Eagle’s Minimum Essential Medium (Gibco, Thermofisher, Dardilly, France) containing 10% FCS. Anti-αvβ3 (LM609) monoclonal antibody directed against human integrin αvβ3 was from Merck Millipore (Darmstadt, Germany). Anti-β1 (MAB17781), anti-β2 (AF1730) and anti-αvβ5 (MAB2528) monoclonal antibodies blocking human integrins β1, β2 and αvβ5 were from Bio-Techne (Lille, France). Mouse isotypic IgG1 antibody (MOPC21) was from ICN Pharmaceuticals (Paris, France). Infliximab was obtained from Dr F. Coury from the Hospices Civils de Lyon (Lyon, France). 1-lysophosphatidylcholine (1-Lyso-PC 18:1), lysophosphatidic acid (LPA, Oleoyl C18:1) were from Interchim (Montluçon, France). Recombinant mouse ATX was generated as a double myc- and His-tag protein and purified as described previously ^29^. Synthetic peptides were purchased from Proteogenix (Schiltigheim, France).

### Cell adhesion assay

Cell adhesion assays were carried out as previously described ^47^. 96-well plates were coated with recombinant mouse, human or rat ATX. Cells were detached and resuspended in HEPES-buffered Tyrode solution supplemented with 2 mM Mn2+ (10^5^ cells in 100 μL of buffer), rested for 1 h at 37°C, and seeded into coated plates for 1 h. Attached cells were fixed, stained with a solution of DAPI. Fluorescent nuclei were counted under the microscope. Results were expressed as specified as the number of attached cells in % of control or in the number of attached cells per well.

### ATX expression and purification

ATX wild type and the various mutants were produced as previously described by Eymery et al (1), using the Flp-In method commercialized by ThermoFisher in order to generate For Generating Constitutive Expression Cell Lines. HEK293-Flp-In cells were cultivated in DMEM with 10% FBS. The cells were grown to 90% confluence, then washed with PBS and typsinized for 10 min with trypsin. Complete media was added to stop the trypsination, the cells were centrifugated 5 min and resuspended in complete media. 10 T300 flasks were used to inoculate 8 roller bottles, that were cultivated for 4 days before switching to a 2% FCS media to make easier the subsequent purification steps. The cells were then left expressing for 4-5 days before collection.

HEK293 medium from 8 roller bottles was combined and the recombinant ATX proteins were purified using a Ni-NTA column preloaded with Ni2+. The column equilibration was performed by washing 10 times with buffer A (20 mM Hepes, 20 mM, 150 mM NaCl, pH 7.4). The protein was eluted using a 5 min linear gradient of buffer B, corresponding to buffer A supplemented with 500 mM Imidazole. Pure fractions were pooled together after SDS-Page analysis. The different fractions were concentrated to 0.5 mL. 5 mg/mL concentrated protein was injected into a Superose 6 (10/300) gel filtration column using buffer A. The purity of the final fractions was analyzed by SDS-Page. The protein concentration was determined by the ratio of the optical density at 260/280 nm using a nanodrop 2000 spectrophotometer (Thermo Fisher Scientific).

### LysoPLD activity

LPC hydrolysis was detected using choline release assay kit in a 96 well plate format as previously described (2). Briefly, 8 μL of amplite red was added to 2 mL of the choline probe before addition of 50 μL per reaction well. The assay was performed by adding 30 nM of recombinant hATX-γ or hATX-β diluted in 50 mM Tris, 150 mM NaCl and 50 μM LPC18:1 or 16:0 at pH 8 to the reaction mixture. A decreasing, 2-fold dilution, of antibodies 31A2 and 35G7 were added in each well starting from 0.1 μM. LPC was added right before the measurement in order to start the enzymatic reaction. Data acquisition was performed using a Clariostar plate reader (BMG Labtech) with fluorescence measurement at λex/λem = 540/590 nm every 60 s for at least 50 min. No inhibition of the reaction was found in the absence of LPC and in the presence of choline, showing a high specificity level of ATX inhibition.

### Western-Blotting and SDS-PAGE

25 μg of hATX WT or hATX mutants’ protein in SDS loading buffer were incubated for 5 min at 95◦ before loading on a 12-well 6–20 % SDS page gel (Invitrogen, # XP04200BOX). The gel was run in a chamber filled with SDS running buffer for approximately 45 min at 225 V. Proteins were detected after staining overnight in Instant-blue.

For western blotting a lower quantity of protein was loaded on a 6–20 % acrylamide SDS-Page gel, 25 to 100 ng, and the gel was not stained. The proteins were transferred onto a nitrocellulose membrane using Trans-blot turbo transfer pack. After transfer, the membrane was blocked in 5 % skim milk powder in TBST for 1 h. Then, the primary antibody was diluted 1 to 1/1000 in TBST with 5 % milk and incubated overnight at 4◦ on the membrane. The blot was then washed three times with TBST and incubated 1 h with a secondary antibody coupled with a peroxidase in 5 % milk in TBST. Before detection, the blot was washed three times with TBST and detection was performed after incubation with the ECL Substrate Kit using the ChemiDoc MP Imaging System (Bio-Rad).

### Column exclusion assay

The proteins were purified by affinity and size exclusion chromatography as previously described. Before injection on a microSEC column Superdex 200 increase 3.2 / 300 (GE healthcare, 28-9909-46), ATX was mixed with the antibody in a 1.2 ratio in presence or absence of 200 μM of LPA for 60 min at 4 degrees. Elution was performed in Tris 50mM, NaCl 150 mM pH 7.4 or in PBS buffer or in Tyrode’s buffer using a microAKTA purifier. The elution was monitored with 280 nm absorbance reading and plotted over time. Futher analysis were conducted by SDS-page analysis of the fraction of interest, as described previously.

### Generation of monoclonal antibodies

The generation of monoclonal antibodies was performed by Biotem company (Apprieu, France) Briefly, specific synthetic polypeptides coupled to KLH were used to immunize mice (OF1 strain, Charles River, Saint Germain de Nuelles, France) subcutaneously in Freund’s complete adjuvant. Lymphocytes were fused to mouse myeloma cells, Sp2/0-Ag14. The antibody-secreting hybridoma cells were selected by a primary screening with an enzyme-linked immunosorbent assay on immunization peptide linked to BSA for anti-β1-IFBS, on both immunization peptide linked to BSA and recombinant mouse ATX for anti-ATX-FLBD antibodies. A competition ATX binding assay of KHOS cells for all antibodies was used as a secondary screening. We established one antibody-secreting hybridoma cell line for anti-β1-IFBS (24E10) and two antibody-secreting hybridoma cell lines for anti-ATX-FLBD (31A2, 35G7). Coding sequences for heavy and light variable chains of antibodies were sequenced and cloned into an IgG2a silent expression vector that was used for transfection of CHO cells allowing the production of recombinant antibodies (rec.35G7), and isotypic control (rec.Isotypic) on a large scale.

### Animal models of collagen-induced arthritis (CIA)

All research involving animals was conducted following institutional guidelines and was approved by the French ethical committee CEEA-55 and the French Ministrère de l’Enseignement Supérieur et de la Recherche (licence no. #34326-2021121018001878). Mice were purchased from Janvier Laboratories (Le Genest, France) and were housed in conventional facilities (ALECS-Conventionnel, Lyon, France). CIA was induced in 8-week-old male DBA1/J mice, as described previously^48^. Mice were immunized by intradermal injection in the tail with an emulsion of type II collagen at 2 mg/ml and complete Freund’s adjuvant (Gentaur, Paris, France) followed by a booster immunization on day 20 by intradermal injection of type II collagen and incomplete Freund’s adjuvant emulsion in the same concentrations, and were sacrificed on day 30. Mice with CIA were treated with the antibodies (10 mg/kg) or vehicle (physiological serum) by intraperitoneal injection twice weekly from day 20 to day 30. Clinical arthritis in each paw was scored on a scale of 0–4 (0: normal, 1: erythema and swelling of one digit, 2: erythema and swelling of two digits or erythema and swelling of the ankle joint, 3: erythema and swelling of three digits or swelling of two digits and the ankle joint, and 4: erythema and severe swelling of the ankle, foot, and digits with deformity). The scores of the four paws were summed to determine a total arthritis score for each animal.

### Microquantitative computed tomography (micro-CT)

Micro-CT analyses of the distal femur and ankles of arthritic mice were carried out using a micro-CT scanner Skyscan 1176 (Skyscan Inc.). The X-ray excitation voltage was set to 50 kV with a current of 500 mA. A 0.5 mm aluminum filter was used to reduce beam-hardening artifacts. Samples were scanned in 70% ethanol with a voxel size of 9.08 μm. Section images were reconstructed with NRecon software (version 1.6.1.8, Skyscan) and 3D images generated with CTvox software (version 3.3.0 r1403, Skyscan).

### Statistical analysis

Statistics were performed, and graphs were generated, using Prism 10 software (GraphPad). Differences between groups were determined by one-way ANOVA followed by Dunnett’s posttest or two-way ANOVA followed by Turkey’s multiple comparison test using GraphPad Prism software (version 10.3.1). Single comparisons were carried out using two-sided unpaired Mann Whitney Test. Effects achieving 95% confidence interval (i.e., p < 0.05) were interpreted as statistically significant. No statistical methods were used to pre-determine sample sizes, but these are similar to those generally employed in the field.

## Acknowledgements.

This work was supported by grants from the INSERM, the Université Claude Bernard Lyon 1, the ANR grant BoneTAX Grant No. ANR-20-CE14-0036-01O and INSERM Transfert grant (OP). The funders had no role in the design of the study; in the collection, analyses, or interpretation of data. CNC acknowledge the support of the Mexico’s French Embassy and Campus France graduate research fellowship program.

## Author contributions

ME performed protein preparation, enzymatic measurements, biochemical experiments, immunological detections, analyzed data and prepared the figures; CNC performed experiments on mice and µCT analysis; LB and CI performed experiments on mice and tissue sampling; FB performed µCT analysis; PCA provided technical advice; AAMC assisted with manuscript writing and data analysis; IMG provided conceptual advice; RL performed cell adhesion analysis; OP provided funding, conceptual advice, coordinated the study and edited the manuscript.

